# Accurate prediction of transcriptional activity of single missense variants in HIV Tat with deep learning

**DOI:** 10.1101/2023.03.16.532970

**Authors:** Houssemeddine Derbel, CJ Giacoletto, Ronald Benjamin, Gordon Chen, Martin R. Schiller, Qian Liu

## Abstract

Tat is an essential gene for increasing the transcription of all HIV genes, and it affects HIV replication, HIV exit from latency, and AIDS progression. The Tat gene frequently mutates *in vivo* producing variants with diverse activities, contributing to HIV viral heterogeneity, as well as drug-resistant clones. Thus, identifying the transcriptional activities of Tat variants will help to better understand AIDS pathology and treatment. We recently reported the missense mutation landscape of all single amino acid Tat variants. In these experiments, a fraction of double missense alleles exhibited intragenic epistasis. It is too time-consuming and costly to determine a variants’ effect for all double mutant alleles with experiments. Therefore, we propose a combined GigaAssay/Deep learning approach. As a first step for determining activity landscapes for complex variants, we evaluated a deep learning framework using previously reported GigaAssay experiments to predict how transcription activity is affected by Tat variants with single missense substitutions. Our approach achieves a 0.94 Pearson correlation coefficient when comparing experimental to predicted activities. This hybrid approach should be extensible to more complex Tat alleles for better understanding the genetic control of HIV genome transcription.

## 1. Background

Human immunodeficiency virus (HIV) causes acquired immunodeficiency syndrome (AIDS), characterized by a progressive failure of the immune system. It remains an important health problem in the United States with 1,189,700 infected people, 18,489 annual deaths, and an annual medical cost exceeding $50 billion dollars[1]. HIV lacks proofreading of its replicated RNA genome and has a high mutation rate of 1 in 10^4^ bp, with each virion 9 kB genome having about 10 new variants [2]. Furthermore, an active HIV infection in a single individual is estimated to generate approximately 10^11^ virions per day [3]. The combination of high mutation rates with efficient virion generation creates extremely genetically heterogeneous and diverse viral genome population, which is a key consideration for important pathogenic processes such as antiretroviral therapy (ARV) resistance, latency, and strain evolution. After selective pressure from ARV therapy, variant virions with drug-resistant variants may survive and propagate, limiting therapeutic efficacy. Therefore, it is important to understand how HIV evolves both within a person, and in worldwide populations with relevance to AIDS pathogenesis and treatment [4].

Tat is an essential regulatory gene that drastically enhances the efficiency of HIV genome transcription and replication. The absence of Tat may lead to short and abortive viral transcripts, and Tat variants widely affect different viral activities. Therefore, comprehensive investigation of various activities of Tat variants can deepen understanding of AIDS pathology and assist drug design targeting for a broader range of HIV-1 strains.

Studies of variant frequencies, HIV evolution, and small-scale mutagenesis studies have greatly advanced knowledge about ARV drug resistance and how to effectively treat AIDS. Although viral isolates from an infected patient are experimentally tested, and the ability to link specific genetic changes with the functions of viral proteins is limited to largely low-throughput experiments, which slowly and incrementally reveals how the vast variant landscape of in a typical infection impacts HIV replication, viral latency, drug resistance, and AIDS pathogenesis.

Two high throughput approaches are available for estimating variants’ effects: the GigaAssay[5] directly measures a functional readout such as transcription, whereas the alternative multiplexed assay of variant effect (MAVE)s are survival screens [6]–[8]. Activities are determined in a GigaAssay by measuring thousands of reads for approximately a million individually UMI-barcoded variant cDNAs. By comparing populations of cDNAs for each mutant to populations for controls, this approach produces accurate measurement and classification of Tat transcriptional activity with high confidence.

Previous analysis of Tat with the GigaAssay reported transcriptional activities for all 1,615 Tat single and 3,429 double missense variants with a ∼95% accuracy [5]. In summary, 35% of all possible single amino acid variants in Tat are loss-of-function. However, it is currently too time-consuming and costly to conduct GigaAssay experiments on millions of variants needed to complete the Tat double missense mutant landscape. We therefore propose an efficient computational method to combine high-accuracy GigaAssay variant/activity data with deep learning algorithms to precisely predicting variants’ effect on Tat activities. The higher performance demonstrated for single missense mutant activity prediction in our results, indicates that this approach can likely be extended to predict the effect of more complex variants and possibly for other protein activities. Our tool is available at https://github.com/qgenlab/Rep2Mut.

## 2. Results

### 2.1 Overview of the proposed deep learning framework called Rep2Mut

We propose to test a deep learning framework called Rep2Mut, to accurately estimate the transcriptional activity of missense variants. The architecture of the Rep2Mut algorithm is shown in *Figure 1* using the Tat protein (86 amino acids) as an example. The output of Rep2Mut is the predicted effect upon transcriptional activity of Tat variants, and the inputs are the wild type (WT) protein sequence, mutated protein sequence (missense variant), and mutated amino acid positions.

**Figure 1:**
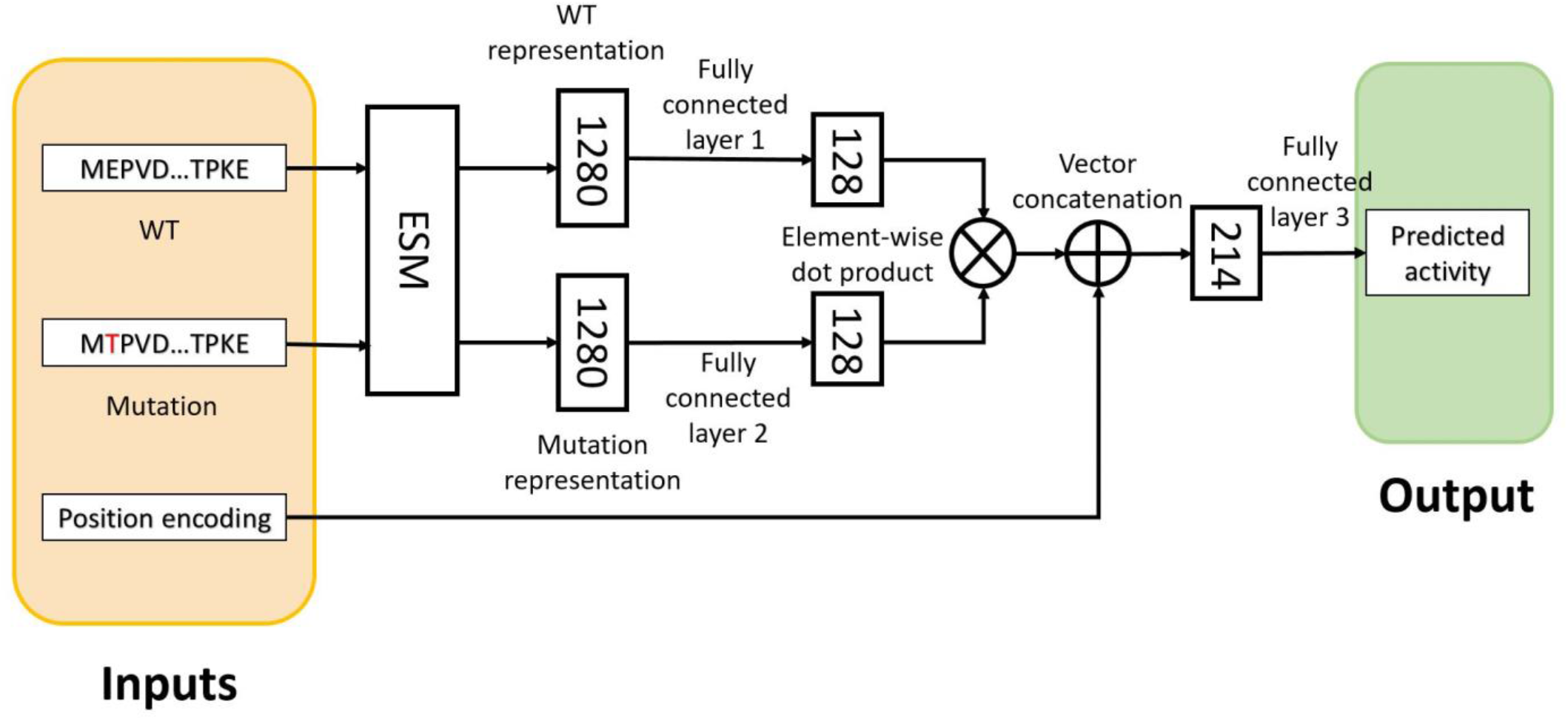
The generic architecture of Rep2Mut model. Tat protein (86 amino acids) is shown as an example. Amino acid in red: mutated residue; Numbers in the rectangles indicates the size of the vectors; filled rectangles in brown: input; filled rectangles in green: output; Cross symbol: elementwise dot product; Plus symbol: concatenation of vectors.

There are several steps in the Rep2Mut algorithm to capture the difference between a WT sequence and its mutated sequences. First, the WT sequence is an input of the evolutionary scale modeling (ESM) protein language model [9] to learn the representation of the position of interest in a WT sequence. This WT representation, with a vector of 1,280 elements, is then used as input of fully connected layer 1 in the network (*Figure 1*) to generate a vector with 128 elements. Similarly, the corresponding mutated sequence is used as input to ESM to generate a representation vector of 1,280 elements in a mutated sequence. The learned representation is then fed into a fully connected layer 2 (*Figure 1*) to generate the other vector with 128 elements. The two vectors of 128 elements are combined by applying element wise multiplication (see *Figure 1*) followed by the concatenation with the position encoding vector of a mutation position. The position encoding vector has N elements (N=86 for the Tat protein); each element is for one position in a protein sequence. All values are zero except for the mutated position which is marked by 1. Next, the combined vector is an input to fully connected layer 3 (*Figure 1*) with a Sigmoid activation function to generate the prediction of transcriptional activity. In total, the Rep2Mut network with the Tat protein has 328,153 trainable weights.

### 2.2 Evaluation of the proposed deep learning framework Rep2Mut

We evaluated Rep2Mut on the GigaAssay transcriptional activity data for all 1,615 single amino acid missense variants in HIV Tat. Layers 1 and 2 of Rep2Mut (in *Figure 1*) were pretrained on 115,997 single variants of 37 existing protein datasets with different protein functional measurements (*Figure 1*). After adding layer 3, all layers in Rep2Mut were then optimized and fine-tuned on the experimental GigaAssay data. To avoid overfitting, 10-fold cross-validation was repeated 10 times and used to calculate the performance of Rep2Mut.

When the variant activities predicted with Rep2Mut were compared to the GigaAssay results, a Pearson correlation coefficient of 0.94 and Spearman correlation coefficient of 0.89 were observed. We repeated the analysis with two recently published methods, ESM [9], DeepSequence [10] and a baseline method, as compared to Rep2Mut in *Figure 2* and *Table 1*. The baseline method is a feed neural network with simple encoding of variant sequences as input (described in Methods), and its performance is ∼0.17 lower than Rep2Mut.

**Table 1:** Pearson and Spearman correlation coefficients comparing experimental activities to predictions from Rep2Mut and state-of-the-art methods.

| Prediction method | Pearson | Spearman |
| --- | --- | --- |
| ESM_pred | 0.51 | 0.56 |
| ESM_pred_avg | 0.54 | 0.59 |
| DeepSequence | 0.57 | 0.41 |
| The baseline method | 0.76 | 0.72 |
| Rep2Mut (wo_p<sup>1</sup>) | 0.91 | 0.87 |
| Rep2Mut | **0.94** | **0.89** |

<sup>1</sup>“wo\_p”: Rep2Mut without position correction vector.

**Figure 2:**
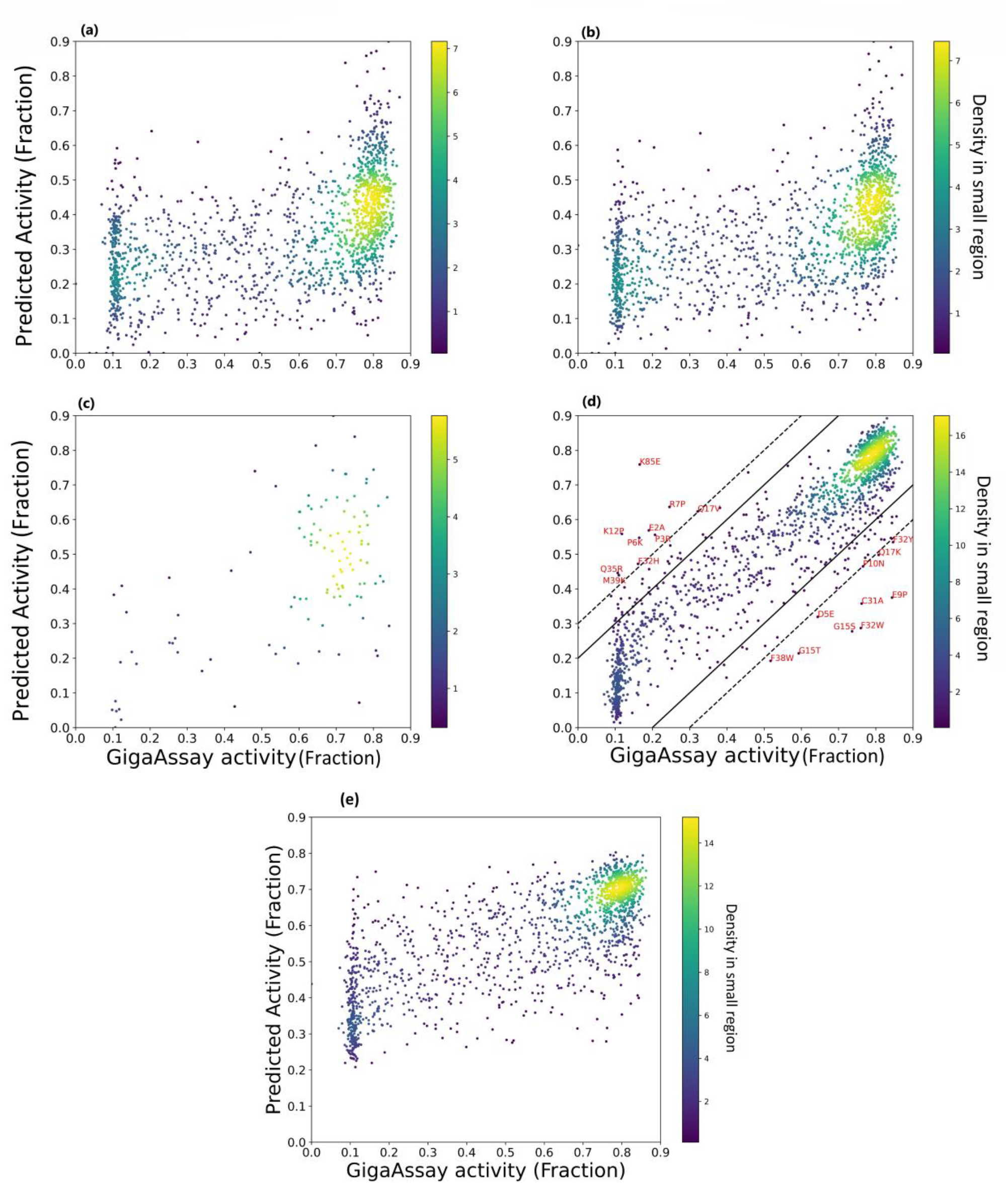
Comparison of activity estimation by Rep2Mut with two state-of-the-art methods. (a) ESM_pred: the best performance among the 5 ESM_pred estimation; (b) ESM_pred _avg: the performance of averaging the 5 ESM_pred estimation; (c) DeepSequence; (d): Rep2Mut; the solid line: the error margins of 0.2 and the dashed line: an error margin of 0.3. Amino acid mutation outliers are labeled with red font. Color legend at the right: the density of the dots in graph; (e) A baseline method.

For ESM prediction methods (called ESM_pred, to be distinguished from ESM models), we tested all Tat variant activities of five trained ESM_pred estimations compared with experimental activities and calculated the best performance. The predictions and the performance of the averaged prediction by ESM_pred is shown in Figure 2 (a) and (b) and in *Table 1*. As expected, the averaged estimation of variants from the five ESM_pred achieves better performance (0.59 Spearman correlation coefficient) than any of the individual ESM_pred estimations.

For the DeepSequence method, we generated multiple sequence alignments using EVision and retrained DeepSequence as suggested by Riesselman et al [10]. DeepSequence needs to be retrained for each protein sequence due to the number of dimensions. DeepSequence generated a higher Pearson correlation coefficient (0.57), but lower Spearman correlation coefficient (0.41) when compared to the ESM_pred prediction of Tat variants’ activities (*Figure 2*(c); *Table 1*). Unfortunately, DeepSequence is only able to generate predictions for 10% of all variant data (see *Figure 2*(c)) even after fine-tuning the retraining process with more sequences in the multiple sequence alignments, demonstrating a limitation of DeepSequence for this application.

In conclusion, our Rep2Mut algorithm achieved much better performance when compared to the state-of-the-art models ESM_pred and DeepSequence models (*Figure 2*(d) and *Table 1*). The Pearson correlation coefficient for Rep2Mut was 0.39 higher than ESM_pred, 0.37 higher than DeepSequence and 0.18 higher than the baseline method. Likewise, the Spearman correlation coefficient was 0.31 higher than ESM_pred, 0.48 higher than DeepSequence and 0.17 higher than the baseline method.

### 2.3 Effect of amino acid position on activity prediction

The activities of Tat variants are partially dependent upon the positions in Tat protein sequence. The last 20 positions of Tat protein (C-terminal) have WT GigaAssay activities with some outliers, such as K85E, which demonstrate a higher tolerance than the N-terminal amino acids (*Figure 3*,). Therefore, we tested the predictions of the Rep2Mut without a position encoding vector. Compared with Rep2Mut, a modified algorithm without a position vector achieves slightly lower Pearson and Spearman correlation coefficients (0.03 and 0.02 lower, respectively; *Table 1*). This result suggests that Rep2Mut has no significant overfitting with positional information.

**Figure 3:**
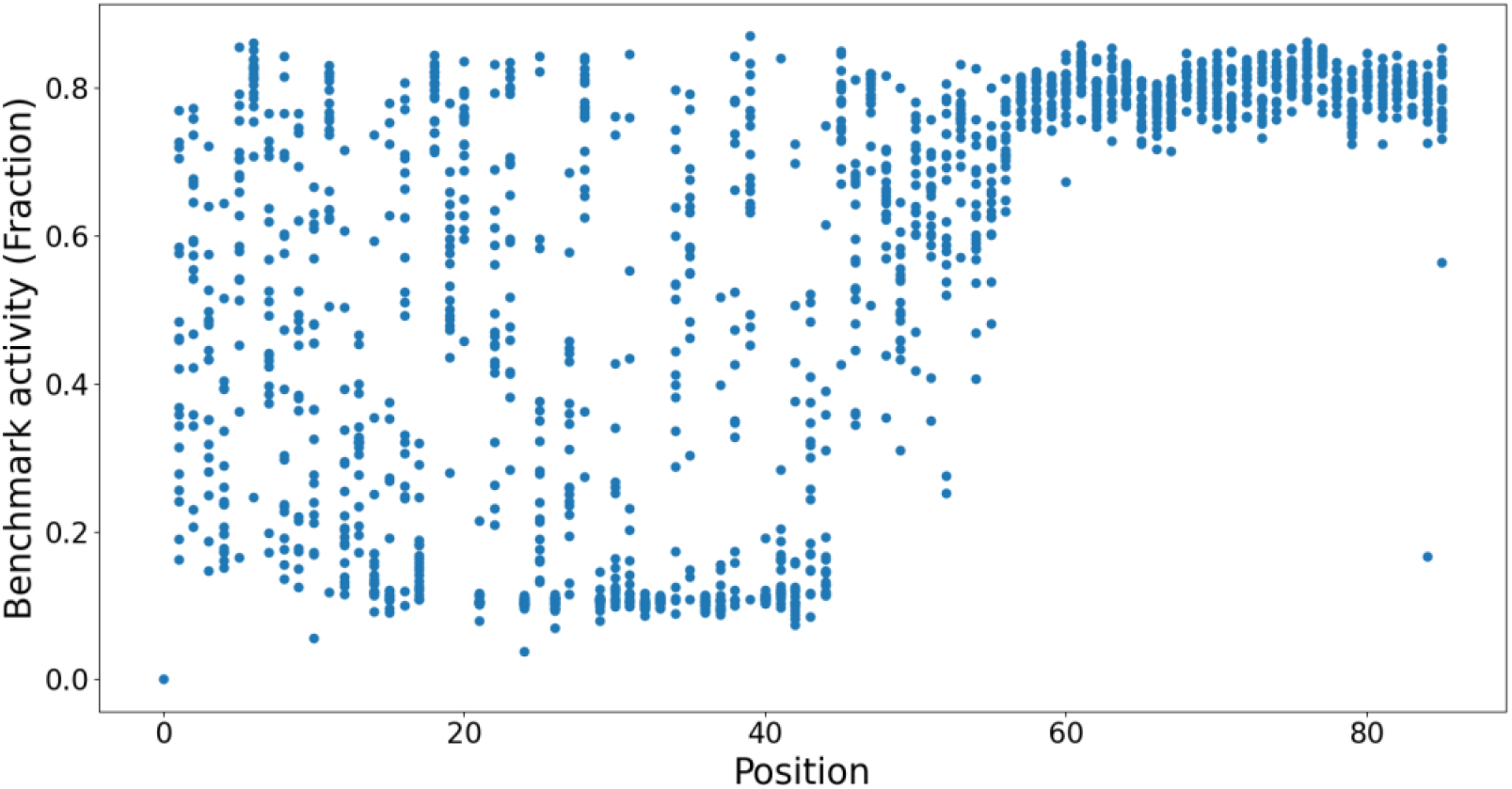
Tat variant activity is partially dependent upon the amino acid position. Each dot represents a Tat variant.

### 2.4 Rep2Mut sensitivity analysis for the fraction of training data

The initial evaluation of Rep2Mut predictions used 90% of the variant activity data (n =1,457) to train Rep2Mut, and the remaining 10% for testing. However, for scaling, wet-lab experiment even with high-throughput approaches such as the GigaAssay, acquiring experimental data is too time consuming and cost prohibitive to assess complex variants. Therefore, we evaluated Rep2Mut’s performance to identify the minimal amount of training data needed to maintain near-maximal performance. We trained Rep2Mut with 70%, 50%, 30%, 20%, 10%, and 7% of the Tat activity single missense variant dataset, and tested Rep2Mut performance with the remainder of the data. To reduce errors from random sampling, we split, trained, and tested those predictions 50 times, calculating an average performance for all tests (*Figure 4*).

**Figure 4.**
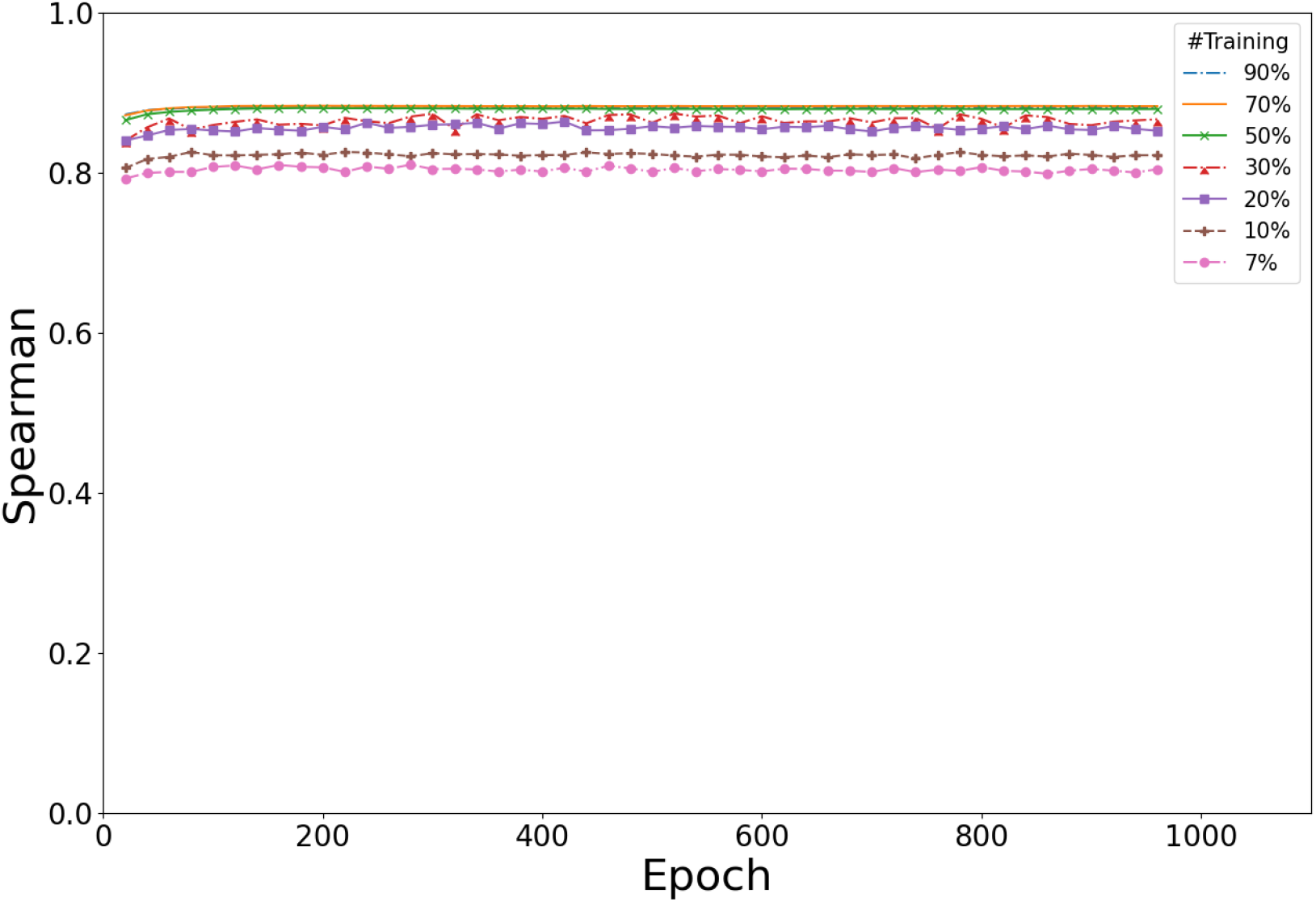
The sensitivity of Rep2Mut performance with the numbers of training instances. X%: X% data are used to train Rep2Mut and (100-X)% for testing, and X is 90, 70, 50, 30, 20, 10, and 7 for different testing strategies.

As expected, reduced Rep2Mut performance was observed with smaller training sampling. However, the performance (Spearman correlation coefficient) had only minimal reduction when Rep2Mut was trained with 50% or more of the data. Further reduction to 20% or 30% of training data, only reduced the performance by 0.03. And the performance further decreases by another ∼0.02 when 10% of data are used for training. Surprisingly, Rep2Mut achieved more than a 0.80 or 0.82 Spearman correlation coefficient when only 7% (113 variants) or 10% of the data, respectively were used for training. This will make the combination of deep learning with the GigaAssay more scalable for multivariant alleles because as little as ∼20% of the variant data can generate predictions of variant effect with little compromise of performance.

## 3. Discussion

### 3.1. Visualization of predicted vectors

To better understand how Rep2Mut predicts variant effects, we used combined vector after dot product in *Figure 1* of all variants to create a global map after dimension deduction to a 2D-space using Uniform Manifold Approximation and Projection (UMAP) [11]. We then investigated the resulting 2-D map for correlations with GigaAssay activity, variant position, and the physiochemical types of amino acids (*Figure 5*).

**Figure 5.**
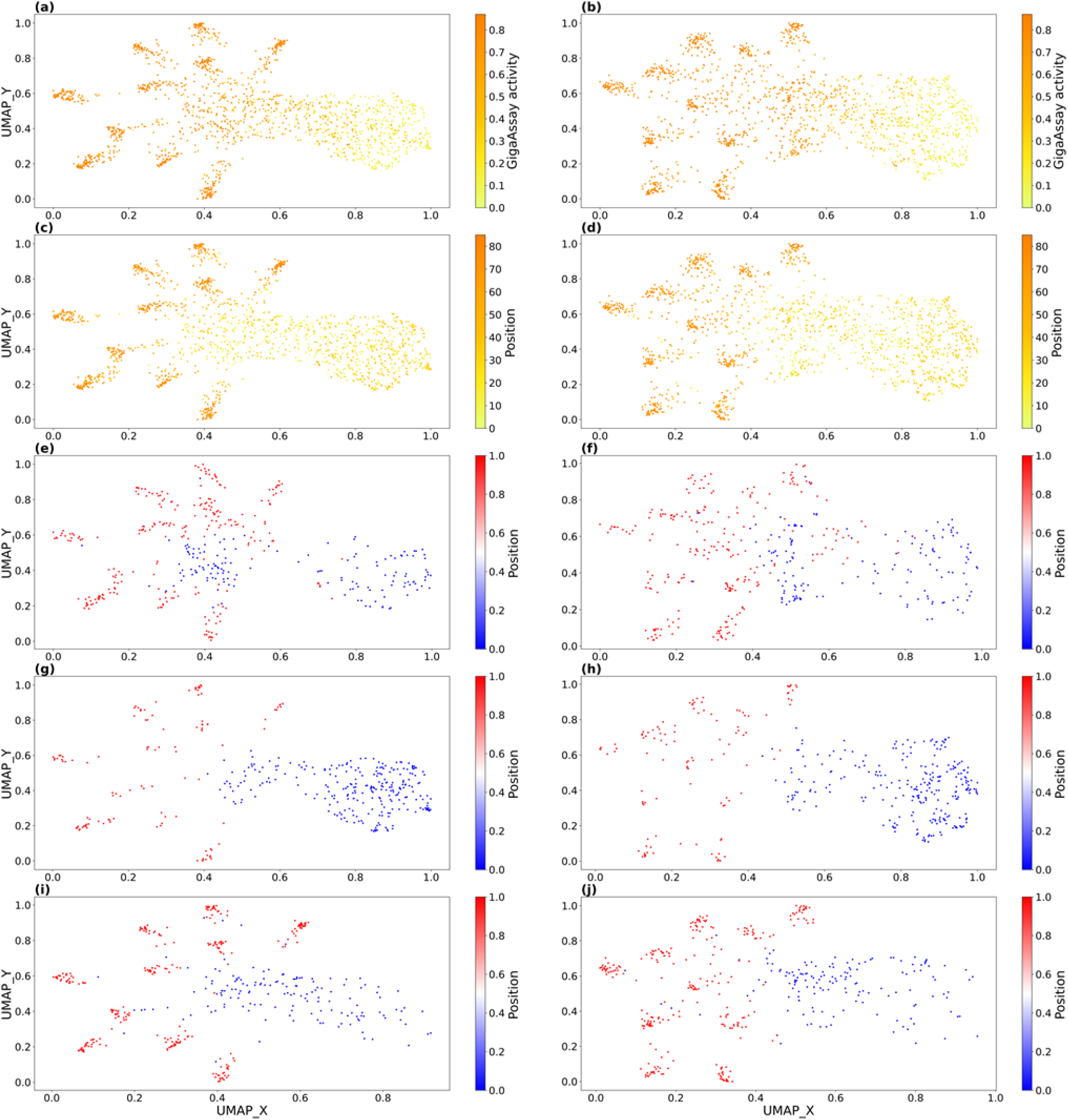
Visualization of Rep2Mut final vectors after dimensionality reduction with UMAP: (a,c,e,g,i): with position vector, (b,d,f,h,j): without position vector, (a,b): colored by GigaAssay activities; (c-j): colored by position; (e,f): positively charged amino acids (Arg, His, and Lys); (g,h): Special cases of amino acids (Cys, Gly, and Pro), (I,j): Polar uncharged amino acid (Ser, Thr, Asn, and Gln). In (e-j), 0: positions of variants lower than 45. 1: positions of variants larger than 45.

*Figure 5* (a) and (b) clearly demonstrate that variants with different experimental activities have a smooth distribution from the right to the left, with or without position encoding. Variants with high activities are in the left half, while most of variants with low experimental activities are in the right half for both *Figure 5* (a) and (b). The positions follow quite similar distribution in the 2D space, although there is abnormal deviation in the middle of the plots of *Figure 5* (c) and (d). This again suggests that the Rep2Mut model itself learned position information from protein sequences without including the position encoding.

### 3.2. Association of amino acid types with Tat activity predictions

*Figure 5* (e-j) show three different types of WT amino acid types in 2D space, with and without the position vector. In all the subplots, there is a clear distribution of experimental activities from right to left. Interestingly, there is no such pattern in the 2D space with different types of mutated amino acids, suggesting non-randomness of WT amino acids at each position.

### 3.3. Outliers in Rep2Mut prediction

To better understand incorrect predictions, we analyzed activity prediction outliers. In *Figure 2*(d) and *Table 2*, we annotated those variants whose predicted activities is 0.3 larger or smaller than the experimentally-determined activities (n = 20). We split them into two groups: overestimation if predicted activities is 0.3 larger than experimental activities, or underestimation if predicted activities is 0.3 smaller than experimental activities. We chose the 0.2 to 0.3 range because this is the approximate error rate for activities determined for UMI barcodes in the GigaAssay [5]. In *Table 2*, we also listed mean, maximum and minimum predicted and GigaAssay activities for the positions of the outlier predictions.

**Table 2.** Overestimated and underestimated outliers predicted by Rep2Mut.

|  |  |  |  | GigaAssay |  |  | Predicted |  |  |
| --- | --- | --- | --- | --- | --- | --- | --- | --- | --- |
| Var | GA | Pred | #ID | Avg | Min | Max | Avg | Min | Max |
| Overestimation |  |  |  |  |  |  |  |  |  |
| E2A | 0.19 | 0.57 | E2 | 0.44 | P=0.16;A=0.19;C=0.24 | D=0.77;T=0.72;Q=0.72 | 0.47 | P=0.28;I=0.36;R=0.36 | S=0.58;T=0.57;Q=0.57 |
| P3R | 0.21 | 0.56 | P3 | 0.56 | R=0.20;K=0.22;G=0.34 | L=0.77;V=0.75;I=0.75 | 0.56 | K=0.41;D=0.43;Y=0.48 | V=0.64;S=0.62;L=0.62 |
| P6K | 0.16 | 0.55 | P6 | 0.62 | K=0.16;R=0.36;L=0.45 | W=0.85;Y=0.79;F=0.77 | 0.61 | R=0.48;D=0.51;E=0.54 | S=0.67;H=0.66;A=0.66 |
| R7P | 0.25 | 0.64 | R7 | 0.78 | P=0.24;K=0.7;I=0.75 | E=0.86;D=0.85;S=0.83 | 0.75 | P=0.63;W=0.66;T=0.69 | E=0.83;Q=0.8;Y=0.8 |
| K12P | 0.12 | 0.56 | K12 | 0.70 | P=0.12;G=0.50;T=0.62 | L=0.83;Q=0.82;N=0.82 | 0.69 | F=0.55;P=0.55;W=0.59 | Q=0.8;A=0.79;T=0.78 |
| Q17V | 0.32 | 0.62 | Q17 | 0.51 | P=0.11;W=0.24;I=0.24 | K=0.8;R=0.78;A=0.77 | 0.51 | P=0.32;F=0.33;Y=0.41 | V=0.62;M=0.61;C=0.59 |
| F32H | 0.16 | 0.47 | F32 | 0.24 | D=0.09;K=0.1;N=0.1 | Y=0.84;W=0.76;L=0.55 | 0.27 | G=0.07;P=0.11;E=0.13 | Y=0.53;H=0.47;M=0.42 |
| Q35R | 0.11 | 0.44 | Q35 | 0.42 | K=0.08;D=0.1;R=0.11 | H=0.79;M=0.74;Y=0.71 | 0.43 | P=0.26;G=0.33;E=0.33 | H=0.55;A=0.55;M=0.51 |
| M39K | 0.11 | 0.45 | M39 | 0.45 | W=0.1;K=0.1;R=0.1 | L=0.84;I=0.78;V=0.78 | 0.45 | P=0.22;R=0.28;D=0.28 | V=0.77;I=0.61;S=0.6 |
| K85E | 0.17 | 0.76 | K85 | 0.76 | E=0.16;W=0.72;F=0.75 | V=0.83;D=0.81;Q=0.81 | 0.78 | P=0.65;M=0.73;W=0.73 | S=0.87;T=0.83;H=0.82 |
| Underestimation |  |  |  |  |  |  |  |  |  |
| D5E | 0.64 | 0.32 | D5 | 0.28 | F=0.15;I=0.16;R=0.17 | E=0.64;S=0.51;C=0.4 | 0.32 | I=0.18;L=0.19;M=0.22 | N=0.46;S=0.46;H=0.41 |
| E9P | 0.84 | 0.37 | E9 | 0.45 | W=0.13;F=0.15;Y=0.17 | P=0.84;A=0.81;D=0.76 | 0.45 | F=0.25;W=0.32;R=0.32 | D=0.66;A=0.61;Q=0.58 |
| P10N | 0.77 | 0.46 | P10 | 0.43 | W=0.12;F=0.14;M=0.17 | N=0.76;A=0.74;S=0.74 | 0.43 | W=0.29;D=0.32;Y=0.33 | A=0.62;S=0.57;C=0.53 |
| G15T | 0.59 | 0.21 | G15 | 0.21 | E=0.09;F=0.11;I=0.11 | S=0.73;T=0.59;P=0.35 | 0.23 | Y=0.11;I=0.14;F=0.16 | Q=0.36;M=0.33;V=0.33 |
| G15S | 0.74 | 0.28 | G15 | 0.21 | E=0.09;F=0.11;I=0.11 | S=0.73;T=0.59;P=0.35 | 0.23 | Y=0.11;I=0.14;F=0.16 | Q=0.36;M=0.33;V=0.33 |
| Q17K | 0.81 | 0.50 | Q17 | 0.51 | P=0.11;W=0.24;I=0.24 | K=0.8;R=0.78;A=0.77 | 0.51 | P=0.32;F=0.33;Y=0.41 | V=0.62;M=0.61;C=0.59 |
| C31A | 0.76 | 0.36 | C31 | 0.24 | P=0.09;Y=0.1;E=0.1 | A=0.76;S=0.73;V=0.42 | 0.25 | R=0.11;K=0.14;D=0.14 | S=0.53;V=0.51;T=0.49 |
| F32Y | 0.85 | 0.53 | F32 | 0.24 | D=0.09;K=0.1;N=0.1 | Y=0.84;W=0.76;L=0.55 | 0.27 | G=0.07;P=0.11;E=0.13 | Y=0.53;H=0.47;M=0.42 |
| F32W | 0.76 | 0.29 | F32 | 0.24 | D=0.09;K=0.1;N=0.1 | Y=0.84;W=0.76;L=0.55 | 0.27 | G=0.07;P=0.11;E=0.13 | Y=0.53;H=0.47;M=0.42 |
| F38W | 0.52 | 0.19 | F38 | 0.15 | D=0.08;Q=0.09;R=0.09 | W=0.51;Y=0.39;L=0.15 | 0.14 | R=0.06;K=0.07;S=0.07 | I=0.22;V=0.22;W=0.19 |

Var: the variants, GA: GigaAssay, Pred: predicted activities, #ID: WT amino acid with variant position, Min: the 3 minimum GigaAssay or predicted values for that variant, Max: the 3 maximum GigaAssay or predicted values for that variant). "X=Y": X is the amino acid type, and Y is the activities.

In all outlier predictions, the average predicted activities of each mutated position are very similar to the averaged experimental activities for that position. The majority of outlier overestimations had very low experimental activities among the 19 variants for each position, while all underestimated outliers had the highest experimental activities. In particular, K12P and K85E, that were overestimated by Rep2Mut, have significantly lower experimental values when compared to other variants at the same positions (K85E: 0.16 vs >0.72 for other K85 variants; K12P: 0.12 vs >0.5 for other K12 variants). Several other variants, such as P3R, P6K, K12P, and R7P, all involve a Proline substitution, suggesting that the unique nature of Proline might not be captured by the deep learning algorithm.

Curiously, all overestimated and underestimated outliers are in the Cyclin T1 interaction site defined in a structure of the Tat:Cyclin T1 complex [12]. Visualization of this structure (PDB: 4OR5) with PyMOL identifies M39K and F32H in two α-helices (27-32 and 34-42) of the Tat protein, and majority of these mutated positions interact with Cyclin T1 (*Figure 6*). According to the accessible surface area calculated by RDBePISA, many of the mutated positions have buried accessible surface area >40 Å^2^ when the Tat protein binding with CyclinT1. The structure analysis also demonstrates that the outliers Q35R, Q17V, and E2A have hydrogen bonds with Cyclin T1. The observation of the erroneous predictions at the Cyclin T1 interface suggests that Cylcin T1 accessibility or interactions may differ in the cell lines used in the GigaAssay. The cells used for the GigaAssay do express Cyclin T1 [13].

**Figure 6.**
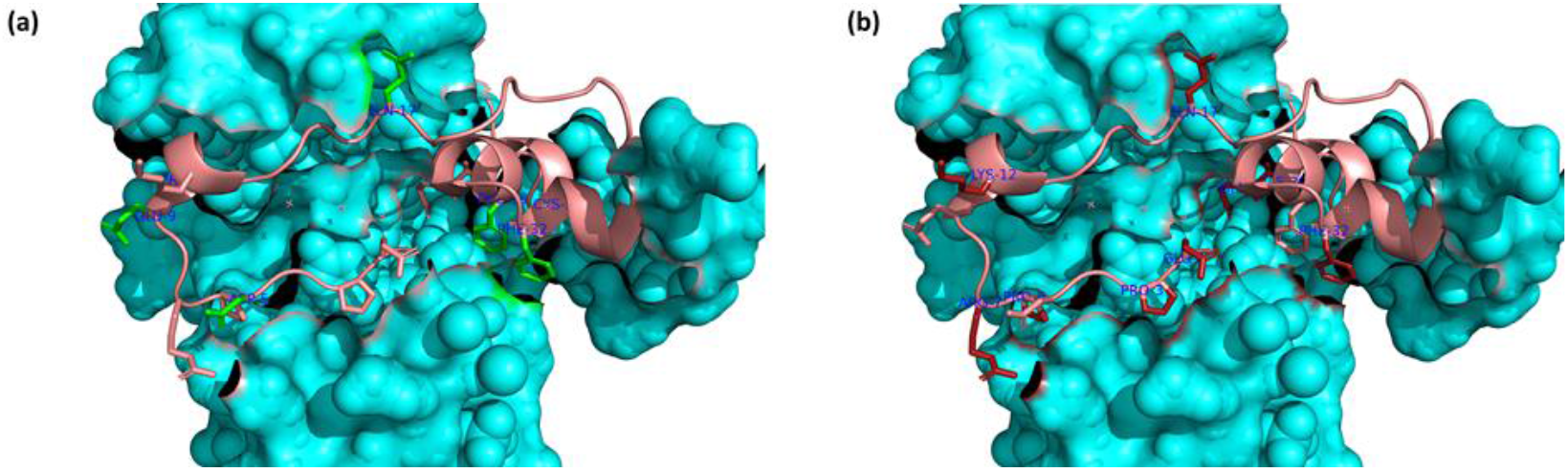
The structure (PDB ID: 4OR5) of the Tat: Cyclin T1 complex. Tat (cartoon) binds to Cyclin T1 (surface view). (a) Underestimated variants are colored green. (b) Overestimated variants are colored red.

## 4. Materials and Methods

### 4.1. Dataset

The dataset used for testing is single variants of HIV Tat proteins, generated by GigaAssay [5]. This Tat protein is composed of 86 amino acids, and each position except the first amino acid is mutated individually to 19 other amino acids besides the WT amino acid. In total, there are 1,615 single missense mutations of Tat protein. Each of missense mutations were sequenced with more than five barcodes, and the transcriptional activity of each variant is calculated by GigaAssay. The effect of single variant is estimated with a value ranging from 0 to 1: the larger the value is, the less effect of the single variant on the transcriptional activities of Tat protein.

### 4.2. Rep2Mut framework to estimate Tat variants’ effect on transcriptional activities

Rep2Mut is a sequence-based prediction of the effect of variants on transcriptional activities measured by GigaAssay. As shown in *Figure 1*, the input of Rep2Mut includes three types of sequence information. One is the WT sequence, and the other is the mutated sequence with a substitution of an amino acid at a position of interest. For either WT or mutated sequence, we used Evolutionary Scale Modeling (ESM) [9] to learn the representation of the mutated position.

ESM [9], [14], [14] is a self-supervised learning framework that was trained on millions of protein sequences to learn multiple levels of protein knowledge from biochemical properties to evolutionary information. It is composed of multiple transformer layers and trained using the masked language modeling objective [15]. Usually, the learned representation at the 33rd layer is used for predicting diverse functions of proteins. ESM-1v [9] is a 34-layer Transformers trained on UniRef90 dataset [16] with 5 released pretrained models. We used the first pretrained model, and fed WT or mutated sequences to it. We used the learned vector of the position of interest at 33^rd^ layer to represent WT or variant information. This learned vector has 1,280 elements.

Each of learned representation vectors from ESM-1v is used as input of a fully connected neural network layer with a vector of 128 elements as output (as shown in *Figure 1*). The PReLU activation function [17] is applied to the layers with a dropout rate of 0.2 to avoid overfitting [18]. The two 128-deminsion vectors are then merged with an elementwise dot product. The element-wise product (or the Hadamard product) is a binary operation that takes two matrices of the same dimensions as input and produces another matrix of the same dimension as the operands. In other words, given two matrices A_m,n_ and B_m,n_, of the same dimension m × n, the elementwise product *A* ⊙ *B* = (*A*)_*ij*_(*B*)_*ij*_ where 0 < *i* ≤ *m*, and 0 < *j* ≤ *n*.

The third type of input to Rep2Mut is a mutated position. A position is encoded into a binary vector of N elements each of which is corresponding to a position in the protein sequence of interest. This encoding vector only has one value of 1 at the mutated positions for a variant and 0 for all other positions. This position encoding vector is then concatenated with the dot-product vector and used as input of another fully connected neural network to predict transcriptional activity. The prediction is normalized with a sigmoid activation function so that the output value ranges from 0 to 1.

### 4.3. Training and testing Rep2Mut

There are two steps to train Rep2Mut. First, we pretrained the layers 1 and 2 (as shown in *Figure 1*) on another 37 protein datasets with various measurements of protein functions. The pretraining is used to optimize weights in the two neural networks. After that, we added layer 3 in *Figure 1*, and fine-tuned Rep2Mut for predicting GigaAssay activities. In both pretraining and fine-tuning processes, we used Adam optimizer [19] and MSE loss function in back-propagation. MSE is defined in Equation *(1)* where n is the number of data points, Y_i_ are the observed activities and Ŷ are the predicted activities.

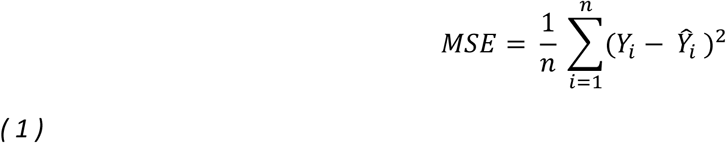

During fine-tuning process, we used a batch size of 8, and a learning rate of 1e-3 for layer 3. Since layers 1 and 2 have been optimized during pretraining step, we used a smaller rate and the learning rate for layers 1 and 2 is 1e-5.

To compare Rep2Mut with other methods, we used 10-fold cross validation. We randomly split the GigaAssay data into 10 groups each of which has 10% of the experimental data. Each time, a group is used for testing and the remaining data for training. We repeated this process ten times and got the mean of performance to evaluate Rep2Mut.

To test the performance of Rep2Mut with different sizes of training data, the variants in the dataset were shuffled, and then split into two sets called training (90%) and test (10%) set. Rep2Mut was learned on training data and evaluated on test data using Pearson and Spearman’s correlation coefficients defined below. To avoid random split, the process above was repeated 50 times, and the averaged performance was calculated for final evaluation.

### 4.4. Evaluation measurements

We used Pearson and Spearman’s correlation coefficients to measure the performance of each tested method. We used the python package scipy to calculate both Pearson and Spearman correlations for the prediction activities of a method. In detail, let X be the GigaAssay activities of a list of variants, and Y be the predicted activities of the same list, and then Pearson correlation coefficients (PCC) are calculated using Equation *(2)* where p is the Pearson correlation coefficient, xi is the ith observed values in X, 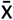 is the mean of X, yi is the ith predicted values in Y and 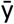 is the mean of Y.

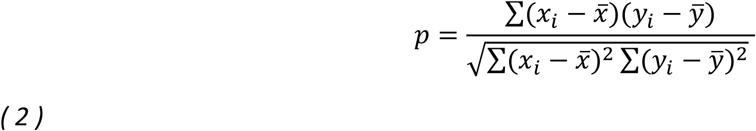

Likewise, the Spearman’s rank correlation coefficients (sPCC) are estimated with Equation *(3)* where sp is Spearman’s rank correlation coefficient, *R*(*) is the ranking of items in *, *cov*(*R*(*X*), *R*(*Y*)) is the covariance of *X* and *Y, σ*_*R*(*X*)_ is the standard deviations of *X*, and *σ*_*R*(*Y*)_ is the standard deviations of *Y*.

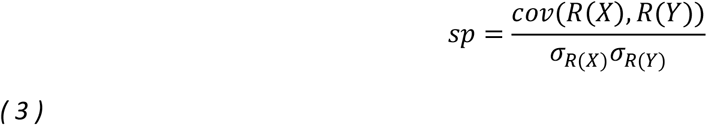

### 4.5. How to use ESM to predict Tat variants’ activities

ESM [9] has diverse capability to estimate proteins’ activities and functions. Here, we used ESM (called ESM_pred so that it is different from ESM released models) to estimate GigaAssay activities that is not done before. For determining variants’ effect, the probability of each amino acids type of a position of interest is predicted in ESM_pred, and the variant effect is calculated based on the logarithmic ratio of the probability between the mutated amino acid and the WT amino acid in Equation *(4)* where T is the set of mutated positions, *x*_\*T*_ is the masked input sequence, 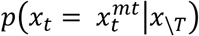 is the probability assigned to the mutated amino acid 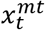, and 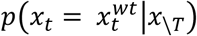 is the probability assigned to the wildtype.

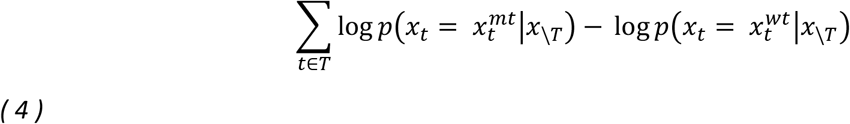

Recommended by ESM [9], five released ESM models (ESM-2 Public Release v1.0.3: esm1v_t33_650M_UR90S_1, esm1v_t33_650M_UR90S_2, esm1v_t33_650M_UR90S_3, esm1v_t33_650M_UR90S_4, and esm1v_t33_650M_UR90S_5) were used individually to predict the transcriptional effect after Tat variants. sPCC was then calculated for each model. In addition, the average prediction for the transcriptional effect of each Tat variant was determined by combining the predictions of five models, and estimated using sPCC.

### 4.6. How to test DeepSequence on Tat variants

DeepSequence [10] is a generative, unsupervised latent variable model for estimating variants’ effect of biological sequences across a variety of datasets with deep mutational scanning. The model was learned in an unsupervised manner solely from sequence information, and grounded with biologically motivated priors, revealing latent organization of sequence families. There are three steps to run DeepSequence. First, a protein sequence of interest was used as input of multiple sequence alignment (MSA) tools to generate multiple sequence alignments. We use recommended tools by DeepSequence, EVcoupling from the website v2.evcouplings.org. DeepSequence [10] suggests the use of a bit score of 0.5 bits/residue as a threshold to generate MSA results. However, MSA results of Tat protein with this score generated only 1.7 Seqs/L with 123 sequences, which is not enough to train DeepSequence. We thus tested 2 bit scores: 0.3 bits/residue with 1,645 effective sequences and 23.8 Seqs/L, and 0.25 bits/residue with 7,871 effective sequences and 110.9 Seqs/L. Second, DeepSequence was trained with the sequences from MSA. Although DeepSequence is a generative model, each protein sequence requires a different model. We used MSA sequences with each bit score to retrain DeepSequence to generate separate models. Finally, the retrained models were used to predict variants’ effect. However, both models predicted only ∼10% (114) of single variants, although 0.3 bits/residue produces better results.

### 4.7. The framework of a baseline method

A simple baseline method was also designed and compared with Rep2Mut. This method uses the one-hot encoding of amino acids at each position as input, and has 3 fully connected layers of feed forward network: the input is a vector of 1,720 elements, the first hidden layer generates a vector of 860, and the second generates a vector of 256. The output is the predicted activity for a variant. This method was trained with a batch size of 16 as well as a learning rate of 5e-4, and tested with the similar strategy as Rep2Mut does.

## 5. Conclusions

We designed a deep learning-based method that only uses protein sequences to accurately predict transcriptional activities of experimentally-determined Tat. With the representation learning with protein sequence models, our approach achieved 0.94 Pearson correlation coefficient. This demonstrates that deep learning-based method can precisely estimate transcriptional activities of proteins with various variants and has great potential to be extended to complex mutations and other protein sequences. Although we use supervised learning, while state-of-the-art methods such as ESM and DeepSequence models use unsupervised training, the superior performance makes our approach more promising for new applications. In particular, our method trained on as little as 20% or 30% of data is able to achieve much better performance than state-of-the-art methods, demonstrating its potential application on other proteins with limited training data. We plan to extend our methods to complex variant alleles and to other proteins for human disease studies.

## References

[1] “Basic Statistics | HIV Basics | HIV/AIDS | CDC,” Apr. 18, 2022. https://www.cdc.gov/hiv/basics/statistics.html (accessed May 06, 2022).

[2] Preston Bradley D., Poiesz Bernard J., and Loeb Lawrence A., “Fidelity of HIV-1 Reverse Transcriptase,” Science, vol. 242, no. 4882, pp. 1168–1171, Nov. 1988, doi: 10.1126/science.2460924.

[3] S. Palmer et al., “Multiple, linked human immunodeficiency virus type 1 drug resistance mutations in treatment-experienced patients are missed by standard genotype analysis,” J Clin Microbiol, vol. 43, no. 1, pp. 406–413, Jan. 2005, doi: 10.1128/JCM.43.1.406-413.2005.

[4] Z. Woodman and C. Williamson, “HIV molecular epidemiology: transmission and adaptation to human populations,” Current Opinion in HIV and AIDS, vol. 4, no. 4, 2009, [Online]. Available: https://journals.lww.com/co-hivandaids/Fulltext/2009/07000/HIV_molecular_epidemiologytransmission_and.5.aspx

[5] R. Benjamin et al., “GigaAssay – An adaptable high-throughput saturation mutagenesis assay platform,” Genomics, p. 110439, Jul. 2022, doi: 10.1016/j.ygeno.2022.110439.

[6] J. Weile and F. P. Roth, “Multiplexed assays of variant effects contribute to a growing genotype–phenotype atlas,” Human Genetics, vol. 137, no. 9, pp. 665–678, 2018, doi: 10.1007/s00439-018-1916-x.

[7] D. Kuang et al., “Prioritizing genes for systematic variant effect mapping,” Bioinformatics, vol. 36, no. 22–23, pp. 5448–5455, 2021, doi: 10.1093/bioinformatics/btaa1008.

[8] L. M. Starita et al., “Variant Interpretation: Functional Assays to the Rescue,” American Journal of Human Genetics. 2017. doi: 10.1016/j.ajhg.2017.07.014.

[9] J. Meier, R. Rao, R. Verkuil, J. Liu, T. Sercu, and A. Rives, “Language models enable zero-shot prediction of the effects of mutations on protein function,” Synthetic Biology, preprint, Jul. 2021. doi: 10.1101/2021.07.09.450648.

[10] A. J. Riesselman, J. B. Ingraham, and D. S. Marks, “Deep generative models of genetic variation capture the effects of mutations,” Nat Methods, vol. 15, no. 10, pp. 816–822, Oct. 2018, doi: 10.1038/s41592-018-0138-4.

[11] L. McInnes, J. Healy, and J. Melville, “UMAP: Uniform Manifold Approximation and Projection for Dimension Reduction.” arXiv, Sep. 17, 2020. Accessed: Jan. 26, 2023. [Online]. Available: http://arxiv.org/abs/1802.03426

[12] J. Gu, N. D. Babayeva, Y. Suwa, A. G. Baranovskiy, D. H. Price, and T. H. Tahirov, “Crystal structure of HIV-1 Tat complexed with human P-TEFb and AFF4,” Cell Cycle, vol. 13, no. 11, pp. 1788–1797, Jun. 2014, doi: 10.4161/cc.28756.

[13] R. Wang et al., “Uncovering BRD4 hyperphosphorylation associated with cellular transformation in NUT midline carcinoma,” Proc. Natl. Acad. Sci. U.S.A., vol. 114, no. 27, Jul. 2017, doi: 10.1073/pnas.1703071114.

[14] Z. Lin et al., “Evolutionary-scale prediction of atomic level protein structure with a language model,” Synthetic Biology, preprint, Jul. 2022. doi: 10.1101/2022.07.20.500902.

[15] J. Devlin, M.-W. Chang, K. Lee, and K. Toutanova, “BERT: Pre-training of Deep Bidirectional Transformers for Language Understanding”.

[16] B. E. Suzek, H. Huang, P. McGarvey, R. Mazumder, and C. H. Wu, “UniRef: comprehensive and non-redundant UniProt reference clusters,” Bioinformatics, vol. 23, no. 10, pp. 1282–1288, May 2007, doi: 10.1093/bioinformatics/btm098.

[17] K. He, X. Zhang, S. Ren, and J. Sun, “Delving Deep into Rectifiers: Surpassing Human-Level Performance on ImageNet Classification,” in 2015 IEEE International Conference on Computer Vision (ICCV), Santiago, Chile, Dec. 2015, pp. 1026–1034. doi: 10.1109/ICCV.2015.123.

[18] N. Srivastava, G. Hinton, A. Krizhevsky, I. Sutskever, and R. Salakhutdinov, “Dropout: A Simple Way to Prevent Neural Networks from Overfitting,” The journal of machine learning research, vol. 15, pp. 1929–1958, 2014.

[19] D. P. Kingma and J. Ba, “Adam: A Method for Stochastic Optimization.” arXiv, Jan. 29, 2017. Accessed: Jan. 26, 2023. [Online]. Available: http://arxiv.org/abs/1412.6980

